# A small molecule PTER-selective inhibitor reduces food intake and body weight

**DOI:** 10.64898/2026.01.26.701829

**Authors:** Sipei Fu, Lushun Wang, Veronica L. Li, Xuchao Lyu, Wei Wei, Xu Shi, Shuliang Deng, Jacob L. Barber, Usman A. Tahir, Charleen Adams, April Carson, Bertha Hidalgo, Laura M. Raffield, James G. Wilson, Hlib Razumkov, Shuke Xiao, Jan Spaas, Daniel Fernandez, Tinghu Zhang, Robert E. Gerszten, Mark D. Benson, Nathanael S. Gray, Stephen M. Hinshaw, Jonathan Z. Long

## Abstract

PTER (phosphotriesterase-related) is an amidohydrolase that mediates catabolism of the anorexigenic taurine metabolite N-acetyltaurine. However, the structural basis of PTER ligand binding and catalysis remain unknown, limiting our ability to harness this pathway therapeutically. Here we solve crystal structures of a eukaryotic PTER in apo and product-bound forms. These structures uncover an unexpected pocket homology between PTER and histone deacetylase (HDAC) enzymes. We exploit this similarity to engineer a first-in-class substrate-competitive PTER inhibitor called PTERi with nanomolar potency and >100-fold selectivity for PTER over HDACs in vitro. Administration of PTERi to diet-induced obese mice reduces feeding, enhances GLP1-RA (glucagon like peptide 1 receptor agonist)-induced weight loss, and prevents weight regain after GLP1-RA discontinuation. The structure of PTER connects histone and metabolite deacetylation into a parallel conceptual framework and enables proof-of-concept data for pharmacological inhibition of PTER in obesity.

## Introduction

Body weight is tightly controlled by genes and biochemical pathways that influence feeding, nutrient metabolism, and energy expenditure. Key to this homeostasis are signals, including metabolites and hormones, that relay information about peripheral nutrient status and energy availability to the brain.^1–3^ Understanding these genetic and biochemical pathways provides fundamental insights into the regulation of energy balance and establishes a foundation for developing therapeutic strategies for the pharmacological manipulation of body weight. For instance, peptides mimetics of incretin hormones such as glucagon-like peptide 1 receptor agonists (GLP-1RAs) are effective anti-obesity therapeutics.^4,5^

Recently, we identified a non-incretin pathway of body weight regulation mediated by the N-acetyltaurine hydrolase PTER (phosphotriesterase-related).^6^ N-acetyltaurine is a conjugate of acetate and taurine whose levels are dynamically elevated upon endurance exercise, dietary taurine supplementation, or other physiologic states associated with increased acetate and/or taurine flux.^7,8^ Catabolism of N-acetyltaurine is mediated by PTER, a body mass index (BMI)-associated amidohydrolase.^6^ Genetic ablation of PTER in mice increases N-acetyltaurine levels, reduces feeding, and confers resistance to diet-induced obesity.^6^ N-acetyltaurine administration is also sufficient to reduce food intake.^6^ These data establish PTER as an enzymatic determinant of N-acetyltaurine levels and a regulator of body weight via its hydrolysis of this anorexigenic metabolite.

From an enzymological perspective, PTER is the only known mammalian N-acetyltaurine amidohydrolase and therefore represents a unique biochemical transformation at the intersection of acetate and taurine metabolism. Nevertheless, the structural basis of PTER catalysis, and how PTER binds ligands, remains unknown. Consequently, our understanding of the PTER/N-acetyltaurine pathway in energy balance remains incomplete, and our ability to harness this pathway therapeutically remains limited. Here, we report the first crystal structures of a eukaryotic PTER in both apo form and in complex with its natural products taurine and acetate. These structures enabled the discovery of convergent pharmacology and pocket homology between PTER and histone deacetylase (HDAC) enzymes.^9^ We exploit this observation to develop a first-in-class selective PTER inhibitor, which reduces feeding, additively enhances GLP-1RA-associated weight loss, and prevents weight rebound following GLP-1RA discontinuation. The structural basis for PTER-dependent N-acetyltaurine hydrolysis therefore provides a unifying conceptual framework for understanding two seemingly disparate modifications – taurine acetylation and histone acetylation – and, in doing so, enables proof-of-concept data for targeting PTER in obesity.

## Results

### Crystal structures of a eukaryotic PTER in apo and product-bound forms

To understand the structural basis for PTER catalysis, we sought to determine crystal structures of PTER in its apo form and in complex with its endogenous taurine and acetate products. To date, experimental structures of eukaryotic PTER proteins have not been reported. We screened >15 eukaryotic PTER orthologs for expression and solubility (**Fig. S1A**). Sponge PTER (sPTER) was selected because of its high protein expression (3 mg/l) and robust N-acetyltaurine hydrolysis activity (**Fig. S1B**). Purified sPTER (>95%) crystallized in the presence or absence of ligands diffracted to high resolution (apo: 2.75 Å; taurine and acetate: 2.4 Å, see **Methods** and **Table S1**). In both cases, crystals grew within three days in similar conditions and had matching space groups and unit cell dimensions, indicating the inclusion of ligands did not affect gross protein or crystal morphology.

sPTER crystallized as a dimer related by non-crystallographic symmetry. The refined structures displayed good geometry and agreement with the data. The resolved sPTER structures had a canonical triose-phosphate isomerase (TIM) barrel fold^10^ consisting of eight alternating α-helices and β-strands (**Fig. 1A,B** and **Fig. S1C**). sPTER also contains a helical insertion region between the first and second α/β modules (sPTER amino acids 35-75), which was conserved among eukaryotic PTER enzymes and commonly absent in bacterial phosphotriesterases (PTEs, **Fig. S1D**). Additional alignment of sPTER with human, rat, and mouse PTERs showed conservation of core secondary structure elements and metal-coordinating residues (**Fig. S1D**). Amino acid side chains lining the substrate access channel and active site were visible in both sPTER structures.

**Figure 1.**
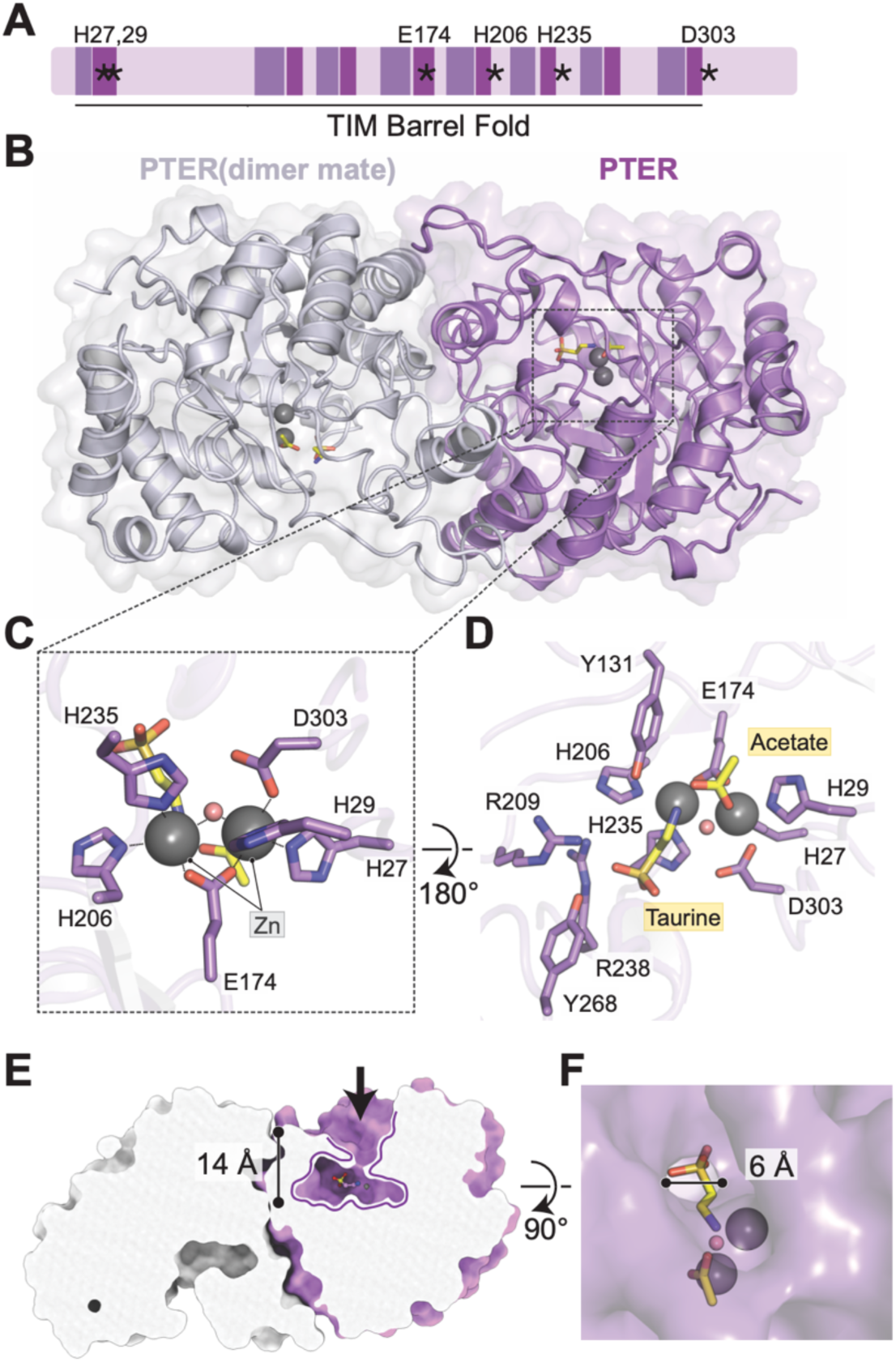
Crystal structure of sPTER in complex with acetate and taurine products. **(A)** Schematic representation of sponge PTER (sPTER) primary amino acid sequence in a linear form (left, N-terminus; right, C-terminus). The domains of the (β/α)_8_-TIM barrel fold sPTER are indicated: β strands in light purple, α helices in dark purple. Asterisks mark the metal-coordinating residues. Black line highlights the TIM barrel fold region. **(B)** Crystal structure of sPTER homodimer in complex with taurine and acetate products. The structure is shown as a cartoon with a semi-transparent surface (grey, chain A; purple, chain B). Products are represented in yellow, with zinc (Zn) ions shown as grey spheres. **(C)** Close-up view of zinc coordination. Metal coordination is shown as black dotted lines. Zn-coordinating water is displayed as a pink sphere. **(D)** Close-up view of taurine- and acetate-interacting amino acid side chains. **(E)** Cutaway view of sPTER surface. The substrate cavity with a depth of 14 Å is outlined as purple curved line. **(F)** Top-down view of sPTER surface with the substrate channel diameter labeled.

A binuclear metal center was located in a deep, central eight-stranded beta barrel channel with clear density defining the positions of both metals (**Fig. 1B,C**). We modeled two Zn atoms into this density. The two metals (Zn-α and Zn-β) were each coordinated by multiple amino acid side chains and bridged by a central water molecule. The more buried Zn-α was coordinated by side chains from His27, His29, Glu174, and Asp303 and water in a trigonal bipyramidal arrangement, while the more solvent exposed Zn-β was coordinated with side chains from Glu174, His206, and His235 and water in a tetrahedral arrangement (**Fig. 1C**).

In the product-bound PTER co-crystal structure, we observed two non-protein electron densities in the PTER active site (**Fig. 1C,D** and **Fig. S2A**). This density could be assigned to acetate and taurine (**Fig. 1C,D** and **Fig. S2A**). Acetate and taurine binding induced only small structural shifts at the PTER dimer interface and minimal changes to the overall structure in comparison to the apo structure (**Fig. S2B**). The acetate ion was situated between the two metals, while taurine density was at a location more distal to the binuclear metal center. The sulfonic group of taurine was oriented facing away from the metal center and the smaller amino group of taurine was pointing towards Zn-β, reflecting its likely position after hydrolysis. The taurine sulfonic group was situated near His206, Arg209, Arg238, and Tyr268, with each oxygen of the sulfonic group forming potential polar interactions at an average distance of 2.9 Å (**Fig. 1D**).

Based on these observations, we propose the following catalytic mechanism: first, N-acetyltaurine traverses the deep substrate access channel and binds near the binuclear metal center. Asp303 then functions as a general base to activate the water bridging the zinc ions. Following hydroxide attack at the N-acetyl carbonyl, Tyr131 functions to stabilize the oxyanion tetrahedral intermediate. Collapse of the tetrahedral intermediate is facilitated by taurine protonation by His305 and subsequent amide bond cleavage (**Fig. S2C**). Consistent with this mechanism, we had previously found that mutations of the orthologous residues in mouse PTER abolished catalytic activity;^6^ our structural data now provide a mechanism by which those residues participate in catalysis.

Lastly, we turned to the question of PTER’s narrow substrate specificity, which we had previously shown to be limited to hydrolysis of N-acetyltaurine alone.^6^ The active site is sufficiently spacious to accommodate larger substrates, and therefore steric hindrance in the active site alone is unlikely to explain the restricted substrate scope (**Fig. 1E**). However, we observed that a long substrate access channel (14 Å in length) provided access to the active site, and that this channel was not uniform in diameter; instead, at its narrowest position the diameter measured only 6 Å (**Fig. 1F**). Four amino acid side chains, Arg209, Phe270, Phe34, and Tyr131, lined the channel at its narrowest position. We therefore propose that this narrow diameter serves as a steric gate, preventing larger substrates from accessing the active site. The presence of polar amino acid side chains in this selectivity filter, and particularly Arg209, likely further restricts the entry of nonpolar metabolites and favors the negatively charged taurine sulfonic acid. Thus, PTER’s substrate specificity appears to arise not from the active site, but rather from a restricted access pathway that selectively permits passage of small substrates.

### Functional homology between PTER and HDAC enzymes

We observed that the deep substrate translocation pathway for N-acetyltaurine to the buried PTER active site (**Fig. 1E,F**) resembles that of the translocation of protein acetyl-lysine side chains into the buried active site of HDACs^11,12^ (**Fig. 2A,B**). Such a deep and narrow translocation pathway is a distinctive feature to both PTER and HDACs^13–16^ that is not shared amongst other deacetylase enzymes for which experimental structures are available.^17,18^ We hypothesized that similar substrate translocation mechanisms might reflect a form of pocket homology of the substrate access channels and active sites between PTER and HDACs. We reasoned that such a functional homology, if present, in principle could be experimentally detected by convergent pharmacology and shared sensitivity to small molecule inhibitors that bind to the corresponding pockets.^19^

**Fig. 2.**
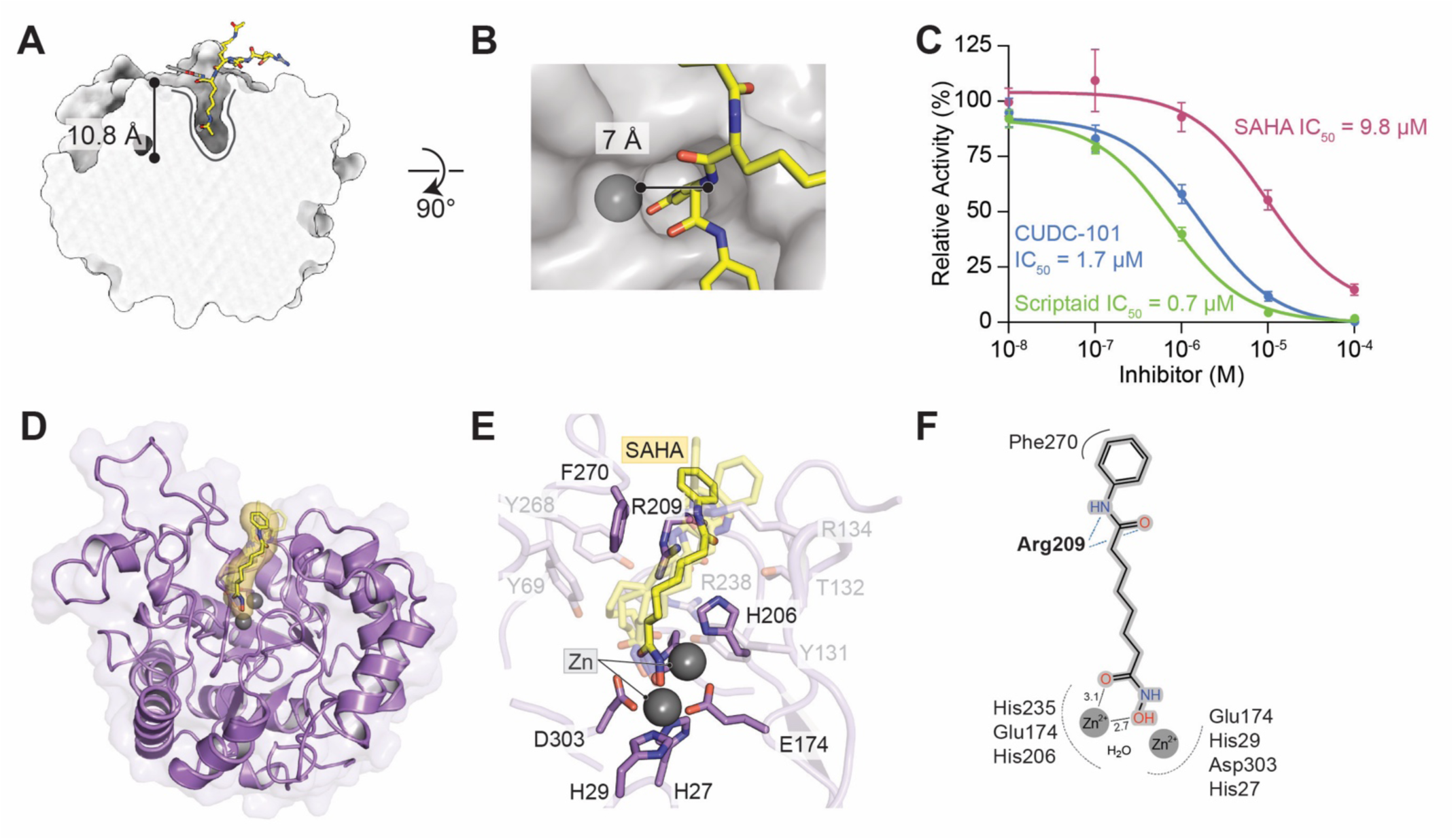
Functional homology and pharmacological convergence between PTER and HDACs. **(A)** Crystal structure of human HDAC8 in complex with a peptide substrate. The structure is shown as a cartoon with a semi-transparent surface. The substrate and its surface are represented in yellow, with zinc (Zn) ion shown as grey sphere. PDB ID: 2V5W. **(B)** Top-down view of HDAC8 surface with substrate channel diameter labeled. **(C)** Dose-response inhibition by the indicated compounds in PTER activity assays. PTER activity (N-acetyltaurine hydrolysis) was measured by quantifying taurine production using 200 ng of purified recombinant mouse PTER (mPTER) and 100 µM N-acetyltaurine for 1 h at 37°C. N=6/data point. All data are shown as mean ± SEM. IC_50_ values for were determined from the dose-response curves via nonlinear regression analysis using GraphPad Prism. **(D)** Top-ranked SAHA pose from molecular docking into the apo-sPTER crystal structure. SAHA and its surface are shown in yellow. **(E)** Top five SAHA poses from molecular docking into the apo-sPTER crystal structure. The top-ranked pose is shown in dark yellow, with the other top poses shown with 30% transparency. SAHA-interacting amino acid side chains are highlighted in dark purple. Zn-coordinating H235 is not labeled for simplicity. Faded protein side chains line the substrate access channel but do not interact with SAHA. **(F)** Schematic representation of interactions between sPTER residues and top-ranked SAHA pose. Metal coordination and hydrogen bonds are shown (grey and blue dotted lines, respectively). Black dotted lines represent average chelation distances (Å) to amide and carbonyl oxygens for the five poses. Hydrophobic interaction is shown in a solid curved line.

To experimentally test this idea, we sought to determine whether SAHA (Vorinostat),^20^ an FDA approved substrate-competitive HDAC inhibitor, can also inhibit PTER. As shown in **Fig. 2C**, SAHA inhibited PTER-dependent N-acetyltaurine hydrolysis with an IC_50_ = 9.8 µM. To further extend these observations, we tested two structurally distinct HDAC inhibitors, CUDC-101 and Scriptaid.^21,22^ These compounds also inhibited both recombinant PTER N-acetyltaurine hydrolysis activity in the low micromolar range (IC_50_ = 0.7 and 1.7 µM, **Fig. 2C**).

We performed molecular docking^23^ of SAHA to understand the potential poses of this compound in PTER. Even in the absence of a pre-defined docking position, all SAHA poses occupied the expected position along the substrate access channel and active site of the PTER protein (**Fig. 2D**). Furthermore, all docked inhibitor poses were highly concordant: the top five models placed the terminal hydroxamate oxygen of SAHA to within ∼4 Å of the less buried Zn-β in the apo-PTER model and the linker regions and the capping groups were in the substrate access channel and near the channel boundary, respectively (**Fig. 2E**). Amino acid side chains from Arg209 and Phe270 were poised to make potential contacts with the distal amide and the hydrophobic capping group, respectively (**Fig. 2E,F**). Therefore, pharmacological convergence between PTER and HDAC in terms of shared inhibitor sensitivity provides experimental confirmation of functional pocket homology. Furthermore, such experiments demonstrate that protein and metabolite deacetylation pathways are connected in a parallel enzymatic framework characterized by sequence-divergent, but pocket homologous deacetylase enzymes.

### Structure-guided engineering of a selective, substrate-competitive PTER inhibitor

Next, we reasoned that a more detailed analysis of the substrate access channels might uncover subtle differences between the two enzymes which could be exploited to develop a PTER-selective inhibitor. By superimposing structures, we observed that all human HDACs feature a pair of conserved aromatic phenylalanine amino acid side chains at a depth of ∼8 Å in the channel (**Fig. 3A**). These residues are positioned to accommodate the hydrophobic alkyl portion of the natural acetyl-lysine side chain substrate. An average spacing of 7.5 Å between these aromatic groups ensures total de-solvation of the alkyl portion of the natural substrate (**Fig. 3A**). SAHA is a substrate-competitive HDAC inhibitor because its hydrophobic alkyl linker mimics the acetyl-lysine side chain (**Fig. 3A,B**). By contrast, the PTER pocket lacks this hydrophobic phenylalanine filter; hydrophilic amino acid side chains are instead found in this location, exemplified by Tyr131, Arg209, and Tyr69 from sPTER (**Fig. 3B**).

**Fig. 3.**
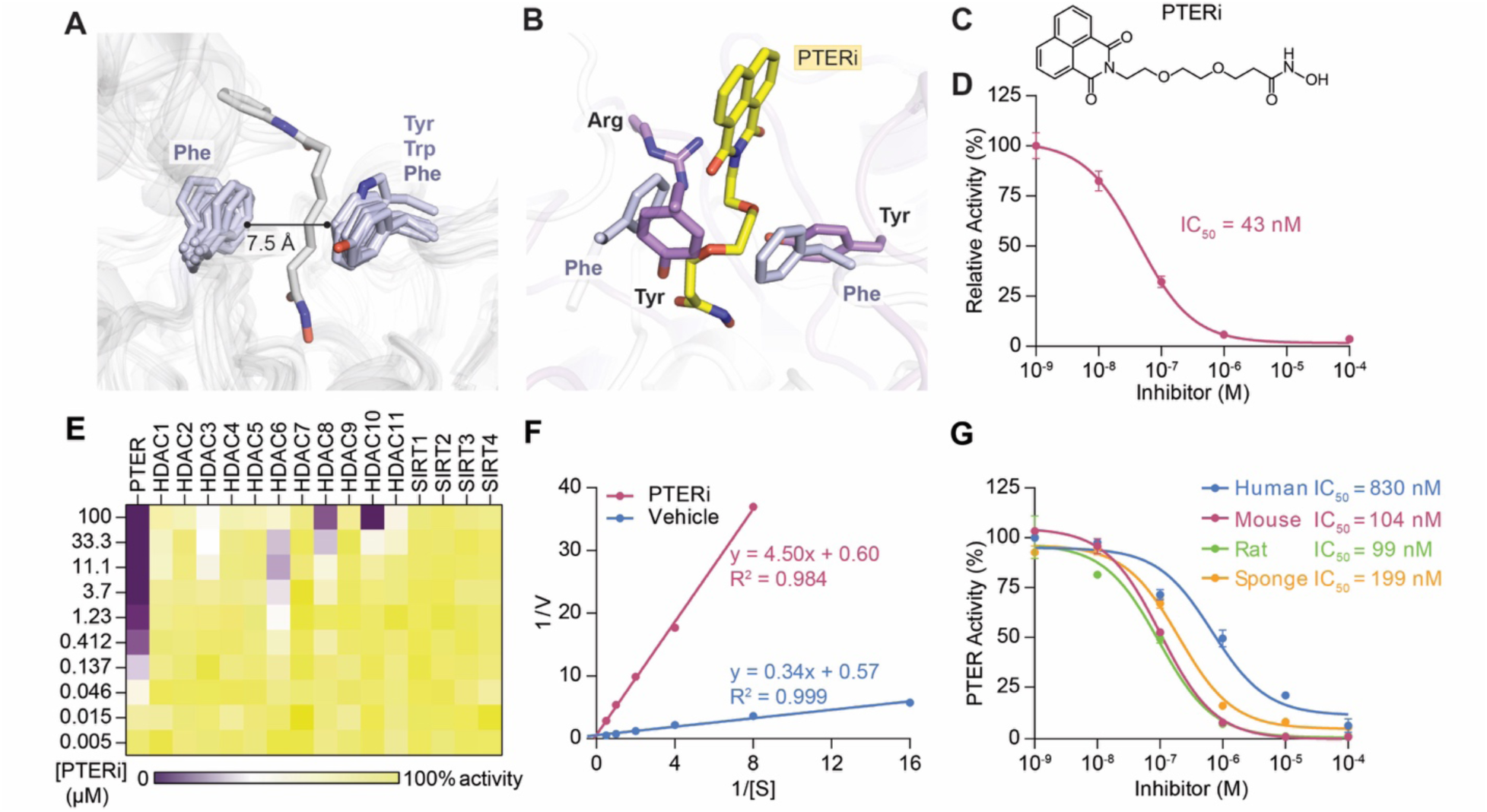
Structure-guided engineering of a PTER-selective inhibitor. **(A)** Superposition of human HDAC(1-11) structures showing conserved aromatic side chains in active site. Average distance between two side chain is labeled. PDB IDs are 4BKX, 4LXZ, 4A69, 2VQJ, 5EDU, 3C0Y, 1T64 for human HDAC 1-4, 6-8. HDAC5 and 9-11 are AlphaFold-predicted models. The representative SAHA (white) is from an HDAC2 co-crystal structure (PDB ID 4LXZ). **(B)** Superposition of the top poses of docked PTERi (yellow) in sPTER and docked SAHA in sPTER, and SAHA from an HDAC2 co-crystal structure (PDB ID 4LXZ). Superposition is based on the top-ranked SAHA pose in sPTER and SAHA pose in HDAC2. SAHA poses are not shown for simplicity. Side chains of sPTER are shown in purple, while side chains of HDAC2 are shown in white. **(C)** Chemical structure of PTERi. **(D)** Dose-response inhibition of PTER activity by PTERi. **(E)** Heat map of dose-response inhibition for PTERi against the indicated recombinant enzyme. **(F, G)** Lineweaver-Burke plot **(F)** and dose-response inhibition of PTERi **(G)** in PTER activity assays. For **(D-G)**, PTER activity (N-acetyltaurine hydrolysis) was measured by quantifying taurine production using 200 ng of purified recombinant mouse PTER (mPTER, panels **D-G**) or purified recombinant PTER from the indicated species **(G)** and 100 µM N-acetyltaurine for 1 h at 37°C. N=3/data point for **(D,G)** and N=1/data point for **(E,F)**. **(D,G)** are shown as mean ± SEM. IC_50_ values for were determined from the dose-response curves via nonlinear regression analysis using GraphPad Prism.

These structural observations suggested that engineering of the appropriate polarity or hydrogen bonding into the linker of SAHA-like compounds might weaken HDAC inhibition by introducing disfavorable interactions with the hydrophobic phenylalanine filter and concurrently increase positive interactions with the analogous hydrophilic amino acid side chains in PTER. The net effect of these minor changes would be magnified by favorable effects on both targets, resulting in improved PTER inhibition and concurrently reduced HDAC activity. After substantial experimentation, we identified an analog of the HDAC inhibitor Scriptaid that accomplishes these design goals (**Fig. 3B,C**). This compound, named PTERi (for “PTER inhibitor”), differs from Scriptaid by the addition of two oxygen atoms into the alkyl linker (**Fig. 3C**). These two oxygen atoms reduce linker hydrophobicity. Molecular docking of PTERi into PTER also shows that the linker oxygens are poised to engage in hydrogen bond interactions with the polar PTER amino acid side chains Tyr131, Arg209, and Tyr69 (**Fig. 3B**). In vitro, PTERi potently inhibited recombinant mouse PTER activity (IC_50_ = 43 nM, **Fig. 3C**). We also examined the selectivity of PTERi for PTER compared to other HDAC enzymes and sirtuin (SIRT) deacetylases. Partial HDAC6 inhibition was observed in the low µM range, while >80% inhibition of HDAC8 and HDAC10 was observed at the highest dose tested (∼100 µM, **Fig. 3E**). Lineweaver-Burke analysis demonstrated that PTERi decreases apparent substrate affinity (e.g., increases K_m_) without affecting maximal enzyme rate (V_max_), indicating that it is a substrate-competitive PTER inhibitor (**Fig. 3E**), as expected based on its expected binding orientation in the PTER pocket and hydroxamate warhead. PTERi also potently inhibited human, rat, and sponge PTER enzymes (**Fig. 3F**). PTERi also inhibited human, rat, and sponge PTER enzymes with IC_50_ concentrations in the nanomolar range (**Fig. 3G**). We conclude that while the overall substrate channel and active site is similar between PTER and HDACs, subtle differences in channel hydrophobicity versus hydrophilicity that have evolved to favor/disfavor their endogenous substrates can be distinguished by appropriately tailored small molecules, enabling the development of a potent and selective PTER inhibitor.

### Acute effects of PTERi on energy balance in mice

The functional homology of PTER and HDAC enzymes now furnished a selective small substrate-competitive molecule PTER inhibitor, which represents a first-in-class chemical tool for pharmacological blockade of PTER activity. All previous studies of PTER in energy balanced have used genetic approaches which produces constitutive and complete elimination of N-acetyltaurine hydrolysis activity;^6^ the availability of PTERi therefore provides a unique opportunity to pharmacologically extend these previous findings.

To determine the effects of PTERi administration in vivo, we treated mice with PTERi (100 mg/kg, IP). This initial dose was selected based on previously reported doses used for SAHA in rodent studies.^24,25^ Plasma PTERi levels peaked at ∼30 µM and exhibited an in vivo half-life *t*_½_ = 9 min (**Fig. S3A**). PTERi was consistently detected in liver and kidney tissues but not brain (**Fig. S3B**). Acute PTERi treatment did not increase liver histone acetylation using two different antibodies (anti-H3K27ac and anti-H4K16ac, **Fig. S3C**). Acute administration of PTERi (100 mg/kg, IP) increased plasma N-acetyltaurine levels by ∼2-fold in both lean (**Fig. 4A**) and diet-induced obese (DIO) mice (**Fig. 4B**). In metabolic chambers, administration of PTERi administration (100 mg/kg, IP) to DIO mice reduced food intake by ∼38% over a 24 h period (P < 0.01, **Fig. 4C**). Overall movement was unaffected (**Fig. 4D**). We also observed a suppression of respiratory exchange ratio, consistent with the reduced food intake and indicative of increased fat oxidation (**Fig. 4E**). Energy expenditure (as measured by VO_2_ and VCO_2_) were unchanged over the 24 h period (**Fig. 4F,G**). The acute anorexigenic effects of PTERi were absent in PTER-KO mice, demonstrating on-target engagement (**Fig. 4H,I**).

**Fig. 4.**
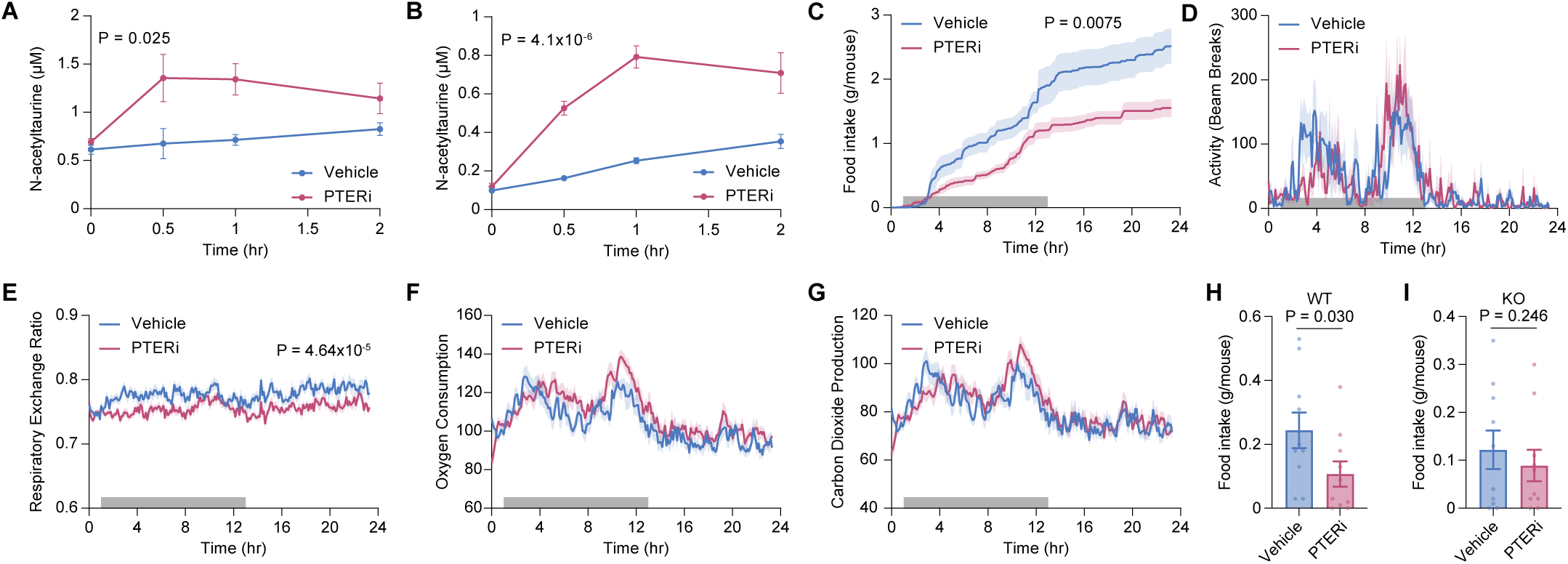
Effects of acute PTERi on energy balance in mice. **(A,B)** Plasma N-acetyltaurine levels after a single administration of vehicle or PTERi (100 mg/kg, IP) to 12-week old male lean mice **(A)** or 22–25-week-old male DIO mice **(B)**. N=5 mice/group. **(C-G)** Cumulative food intake **(C)**, ambulatory activity **(D)**, respiratory exchange ratio **(E)**, oxygen consumption **(F)**, and carbon dioxide production **(G)** of 15-week-old male DIO mice over a 24 hours period after a single acute administration with either vehicle or PTERi (100 mg/kg, IP). Gray bar, night cycle. N=12 mice/group. Initial body weights, mean ± SEM: vehicle, 39.5 ± 1.9 g; PTERi, 41.4 ± 1.3 g, P > 0.05. **(H,I)** Food intake over 3 h after a single administration of vehicle or PTERi (100 mg/kg, IP) to 24-week-old male WT **(H)** or PTER-KO **(I)** mice. All data are shown as mean ± SEM. P-values for **(A-G)** were calculated by two-way ANOVA. P-values for **(H,I)** were calculated by Student’s *t*-test.

Because the anorexigenic effects of exogenous N-acetyltaurine treatment had previously been shown to require functional GFRAL receptors in mice, we examined circulating GDF15 protein and *Gdf15* mRNA levels following PTERi treatment. Plasma GDF15 levels were increased by PTERi (**Fig. S4A**), as were levels of *Gdf15* mRNA in the liver, but not quadriceps, gut, spleen, or kidney (**Fig. S4B**). PTERi also induced a conditioned taste aversion response (**Fig. S4C**). Lastly, the acute anorexigenic effects of PTERi were absent in mice treated with a previously described anti-GFRAL neutralizing antibody (Eli Lilly & Co., antibody clone 8A2,^26^ **Fig. S4D**).

### Chronic effects of PTERi on body weight in mice

To determine if chronic administration of PTERi would durably suppress food intake and body weight, we treated DIO mice daily with PTERi (100 mg/kg/day, IP) or vehicle control. Over a 4-week study period, PTERi-treated animals ate ∼12% less than vehicle-treated mice (**Fig. 5A**). At the end of the experiment, PTERi-treated mice lost -2.2 ± 0.6 g (mean ± SEM) of body weight, whereas vehicle-treated mice gained 1.3 ± 0.5 g (mean ± SEM), producing a 9% net vehicle-adjusted weight loss in PTERi-treated animals (**Fig. 5B**). Dissection of tissues at the end of the experiment revealed a loss of adipose mass with no changes in the masses of the other organs, including liver, kidney, and heart (**Fig. 5C**). Upon study termination, blood liver enzymes aspartate aminotransferase (AST) and alanine aminotransferase (ALT) were not different between treatment groups (**Fig. 5D,E**), demonstrating that chronic PTERi treatment does not produce liver toxicity. Hematoxylin & eosin (H&E) staining of adipose tissues and liver from PTERi-treated mice was did not reveal any obvious differences between treatment groups (**Fig. 5F,G**). Measurement of mRNA levels for a panel of inflammatory genes from adipose and liver tissues also did not reveal any induction of an inflammatory gene expression program in response to PTERi (**Fig. S5A,B**). A broader metabolic panel revealed that chronic PTERi treatment was associated with a ∼35-50% reduction in total and HDL-cholesterol without changes to LDL-cholesterol and a ∼25% reduction of non-fasted blood glucose without changes to insulin levels (**Fig. S5C-G**). We also observed a modest increase in plasma triglycerides without changes to plasma non-esterified free fatty acids (**Fig. S5H,I**). Total liver triglycerides were unchanged in mice chronically receiving PTERi (**Fig. S5J**). Mitochondrial protein content was also unchanged in muscle or liver between PTERi-treated and vehicle-treated groups (**Fig. S5K,L**). Lastly, the effects of PTERi on N-acetyltaurine levels, food intake, and body weight were dose-dependent (**Fig. S5M-O**).

**Fig. 5.**
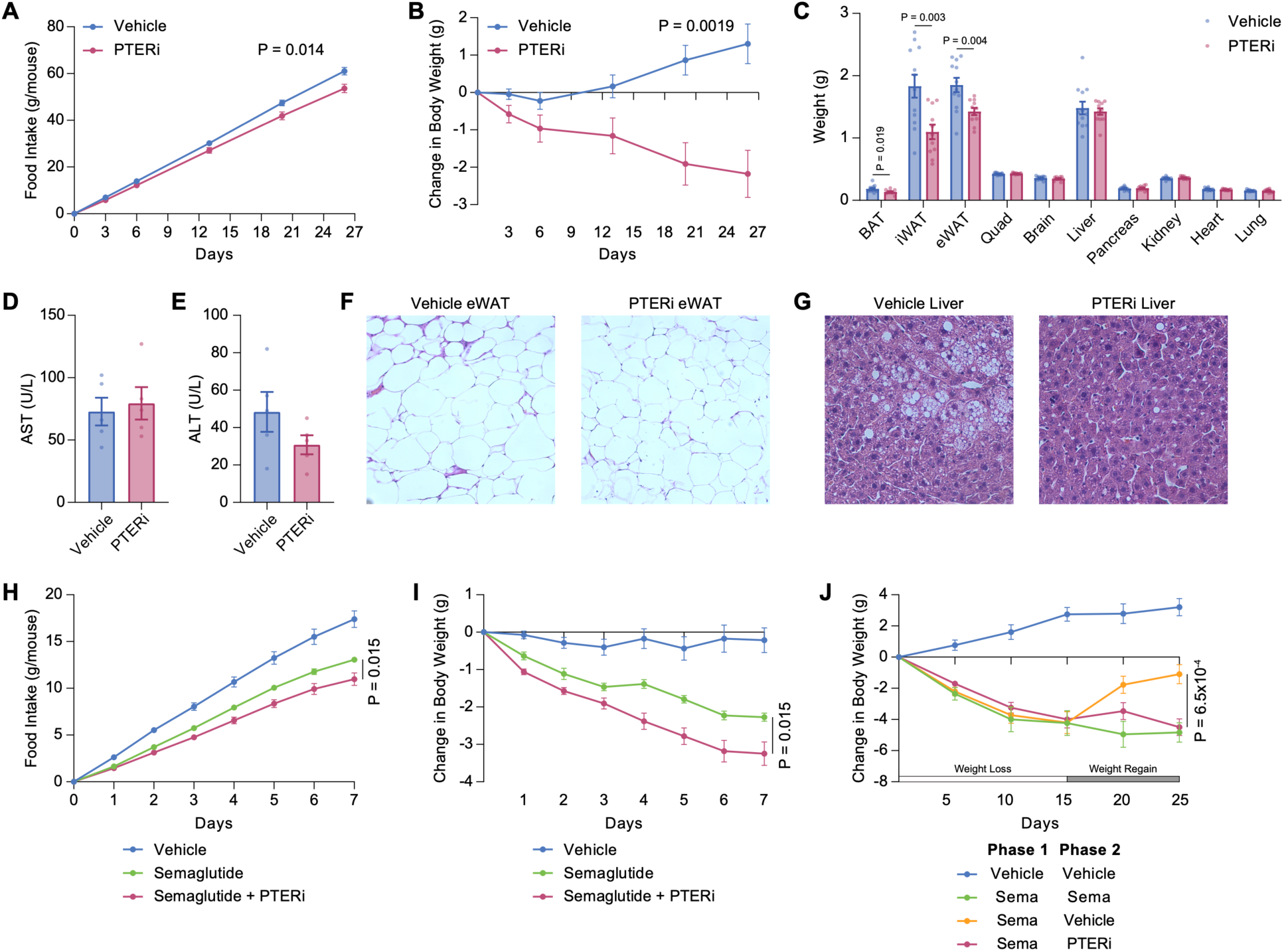
Effects of chronic PTERi on energy balance in mice. **(A-C)** Cumulative food intake **(A)**, change in body weight **(B)**, and tissue weights at the end of the study **(C)** from 18-week-old male DIO mice following chronic administration with vehicle or PTERi (100 mg/kg/day, IP). N=12 mice for vehicle group and N=11 mice for PTERi group. Initial body weights, mean ± SEM: vehicle, 39.3 ± 1.0 g; PTERi, 38.1 ± 0.9 g, P > 0.05. For **(C)**, BAT, brown adipose tissue; iWAT, inguinal white adipose tissue; eWAT, epididymal white adipose tissue; quad, quadriceps. **(D-G)** Aspartate transaminase (AST) **(D)**, alanine transaminase (ALT) **(E)**, eWAT hematoxylin and eosin (H&E) staining **(F)**, and liver H&E staining **(G)** from 18-week-old male DIO mice following chronic administration with vehicle or PTERi (100 mg/kg/day, IP) for 27 days. Initial body weights, mean ± SEM: vehicle, 39.3 ± 1.0 g; PTERi, 38.1 ± 0.9 g, P > 0.05. **(H,I)** Cumulative food intake **(H)** and change in body weight **(I)** of 18-week-old male DIO mice following chronic administration with either vehicle, Semaglutide (0.3 nmol/kg/day, IP), or a combination (Semaglutide 0.3 nmol/kg/day, IP, and PTERi 100 mg/kg/day, IP). N=10 mice/group for vehicle or Semaglutide treated mice. N=9 mice/group for combination (Semaglutide and PTERi). Initial body weights, mean ± SEM: vehicle, 37.2 ± 1.3 g; Semaglutdie alone, 38.1 ± 0.8 g, combination Semaglutide+PTERi, 36.3 ± 0.9 g. **(J)** Change in body weight of 21-week-old male DIO mice following administration with Semaglutide (2 nmol/kg/day, IP) or vehicle during the weight loss phase (horizontal white bar), followed by vehicle, Semaglutide (2 nmol/kg/day, IP), or PTERi (100 mg/kg/day, IP) during the weight regain phase (horizontal grey bar). N=5 mice/group for Vehicle/Vehicle and Sema/Sema; N=10 mice/group for Sema/Vehicle and Sema/PTERi. Initial body weights for all groups, mean ± SEM: 43.9 ± 1.0 g. All data are shown as mean ± SEM. P-values for **(A,B** and **H-J)** were calculated by two-way ANOVA. P-values for **(C)** were calculated by Student’s *t*-test.

To determine the efficacy of PTERi when used directly in combination with GLP-1RAs, DIO mice were treated daily with either the GLP-1R agonist Semaglutide alone (0.3 nmol/kg/day, IP) or a combination of Semaglutide and PTERi (0.3 nmol/kg/day Semaglutide, IP, and 100 mg/kg/day PTERi, IP). Over a one-week treatment period, Semaglutide-treated mice ate 25% less than vehicle-treated mice, resulting in a total weight loss of -2.3 ± 0.1 g (mean ± SEM) compared to vehicle-treated mice (**Fig. 5H,I**). In the Semaglutide and PTERi combination group, cumulative food intake was further reduced (-37% less than vehicle-treated mice), and body weight was correspondingly further lowered compared to vehicle-treated mice (mean ± SEM, -3.2 g ± 0.3 g, P < 0.001, **Fig. 5H,I**). We conclude that PTERi treatment lowers body weight via a mechanism that is additive with GLP-1RAs, as would be the expected for the pharmacological targeting of two distinct molecular pathways that regulate energy balance.

Next, we sought to test PTERi in a mouse model of incretin discontinuation-associated weight rebound. Here, DIO mice were treated with Semaglutide (2 nmol/kg/day, IP) for 15 days (weight loss, phase 1) and then animals were switched to vehicle for an additional 10 days (weight regain, phase 2). The experimental group consisted of DIO mice treated with Semaglutide (2 nmol/kg/day, IP) during the weight loss phase and then switched to PTERi (100 mg/kg/day, IP) during the weight regain phase. As additional comparators, control groups of DIO mice were maintained on only vehicle or only Semaglutide (2 nmol/kg/day, IP) for the entire treatment protocol. During the initial treatment phase, all mice lost body weight (mean ± SEM, -4.1 g ± 0.4 g) whereas vehicle-treated mice gained weight (mean ± SEM, +2.7 g ± 0.4 g, P < 0.001, **Fig. 5J**), resulting in a ∼16% net vehicle-adjusted weight loss. In the weight regain phase, Semaglutide-treated mice that were switched to vehicle treatment rapidly gained weight (mean ± SEM, +3.1 g ± 0.5 g). By contrast, Semaglutide-treated mice that were then switched to PTERi maintained their weight loss (mean ± SEM, -0.5 g ± 0.7 g, P < 0.001) at a level comparable with mice that continued Semaglutide therapy (mean ± SEM, -0.6 g ± 0.3 g, **Fig. 5J**). We conclude that PTERi is effective for weight maintenance and the prevention of weight regain following GLP-1RA discontinuation.

To correlate the metabolic effects of PTERi treatment with the biochemical activity of this compound, we synthesized an inactive analog of PTERi (PTERamide) by chemical replacement of the hydroxamate with a primary amide. This modification eliminates the metal chelation activity of the PTERi hydroxamate group and would therefore be expected to abolish its ability to block PTER enzyme activity (**Fig. S6A**). Using recombinant PTER protein and N-acetyltaurine hydrolysis activity assays, we observed that PTERamide was, as expected, completely ineffective in blocking PTER enzyme activity even at 100 µM (**Fig. S6B**). Likewise, PTERamide (100 mg/kg, IP) did not reduce acute food intake (**Fig. S6C**) and did not lower body weight upon chronic administration (100 mg/kg/day, IP, **Fig. S6D**).

## Discussion

PTER (phosphotriesterase-related) is a recently de-orphanized amidohydrolase that mediates degradation and inactivation of the anorexigenic taurine metabolite N-acetyltaurine.^6^ By determining the co-crystal structure of a eukaryotic PTER in complex with its acetate and taurine products, here we uncover an unexpected pharmacological convergence and pocket homology between PTER and HDAC enzymes. We leverage this insight to generate a first-in-class PTER-directed small molecule inhibitor and provide evidence that pharmacological inhibition of PTER in mice reduces body weight. Our work connects the regulation of protein and metabolite acetylation through parallel enzymatic processes and enabling proof-of-concept data for pharmacological targeting of PTER in obesity.

PTER and HDAC enzymes are evolutionary distinct in both sequence and overall structure. Nevertheless, both enzymes catalyze deacetylation reactions, and their shared sensitivity to SAHA and related small inhibitors provide direct evidence for pharmacological convergence and a functional pocket homology between their substrate channels and active sites. This unexpected convergence, beyond the practical aspects of enabling a first-in-class PTER inhibitor, also offers broader insights into the role of acetylation as a chemical modification in biology. Classically, histone acetylation acts as an activating post-translational modification that promotes gene expression. Similarly, taurine acetylation can now be interpreted as an “activating” metabolite modification, stimulating a neomorphic action for N-acetyltaurine (anorexigenic activity). In both cases, these neomorphic activities are terminated by their respective deacetylases – HDAC for acetylated histones and PTER for N-acetyltaurine – revealing a unifying chemical logic and enzymatic regulatory paradigm across seemingly disparate post-translational modifications.

Obesity remains a pressing public health concern.^27^ Over the last decade, GLP-1RAs have emerged as a powerful and remarkably safe pharmacological strategy for lowering body weight in humans.^5,28^ Nevertheless, there remains considerable interest in the identification and development of non-GLP-1R strategies for the therapeutic modulation of body weight.^29–31^ Here we introduce PTERi as a tool compound that serves as a starting point for the development of small molecules that block PTER activity for obesity. PTERi already achieves ∼9% weight loss as a single agent and is additive with, and complementary to, GLP-1RAs, as would be expected for pharmacological targeting of a differentiated, non-incretin mechanism.^29^

Several prior studies have shown that blockade of HDACs can drive weight loss and mitigate obesity-related complications in mouse obesity models. For instance, HDAC6 inhibition reduces food intake and body weight via leptin-dependent pathways,^32,33^ while blockade of several other HDACs, including HDAC1,^34^ HDAC3,^35,36^ and HDAC11,^37^ confers resistance to diet-induced obesity via energy expenditure pathways. We confirmed that the acute anorexigenic activity of PTERi is due to on-target PTER engagement using PTER-KO mice. PTERi treatment also did not increase whole-body energy expenditure in metabolic chambers. Therefore, the molecular and physiologic effects of PTERi on energy balance are fundamentally distinct from these previous HDAC-driven mechanisms.

As pharmacological efforts towards PTER progress forward, some caution is also warranted. Side effects of PTER blockade, such as nausea, will also need thorough evaluation, particularly in comparison to GLP-1RAs. Importantly, *PTER* mRNA is highly expressed in microglia and astrocytes of the human cerebral cortex and not specifically enriched in areas of the brain that regulate energy balance. Mood changes and other untoward psychiatric side effects have been observed with anti-obesity agents that act more broadly in the brain, such as the CB1 antagonist Rimonabant.^38^ Because of the need for anti-obesity drugs to be very safe over long-term use, the psychiatric, cardiovascular, and metabolic safety profile following pharmacological blockade of PTER with will need to be very carefully evaluated in future studies.

Addressing these concerns will require a broad and concerted effort towards chemical discovery of improved compounds, which is now enabled by structural information about a eukaryotic PTER enzyme. The development of next generation PTER inhibitors with improved pharmacokinetics, prolong target engagement in vivo, and enhanced selectivity will enable the long-term efficacy, tolerability, and safety of small molecule PTER blockade to be thoroughly evaluated in humans.

### Limitations of Study

There are four limitations of this study. First, although our crystal structure represents the first eukaryotic PTER (derived from sponge), it may not capture mammalian-specific details in the enzyme’s binding site or catalytic function. Future structural work on mammalian PTER could reveal important nuances relevant to therapeutic targeting. Second, we use computational modeling to predict how inhibitors fit into the PTER active site. In the future, co-crystal structures with inhibitors would help clarify inhibitor binding modes and refine medicinal chemistry efforts. Third, PTERi itself is best viewed as a research tool rather than a drug candidate owing to its short in vivo half-life and potential off-target effects. With respect to potential off-target effects, our PTERi compound exhibits IC_50_ ∼50 nM for PTER. Partial HDAC6 inhibition was observed in the micromolar range, while >80% inhibition of HDAC8 and HDAC10 was observed at the highest dose tested (∼100 µM). In the future, rigorously determining the potential contribution of other HDACs to the effects of PTERi will require testing the effects of this compound in the corresponding HDAC knockout mice. Alternatively, PTER-KO mice and anti-GFRAL antibodies could also be useful for fully delineating potential PTERi on-target versus off-target activity in the chronic setting. Fourth, while we show proof-of-concept efficacy of PTERi in phenotypes related to energy balance, we do not directly examine any other of the phenotypes that would be necessary to establish safety, such as psychiatric and neurocognitive side effects, cardiovascular safety, renal safety, or drug interactions with other commonly used medications in obesity.

## RESOURCE AVAILABILITY

### Lead contact

Further information and requests for reagents may be directed to and will be fulfilled by the lead contact Dr. Jonathan Z. Long.

### Materials availability

Further information and requests for reagents will be fulfilled by Dr. Jonathan Z. Long. A list of critical reagents (key resources) is included in the key resources table. Material that can be shared will be released via a Material Transfer Agreement.

### Data and code availability

All data generated or analyzed during this study are included in this published article and its supplementary information files. No new code was generated in this study.

## ACKNOWLEDGMENTS

We thank members of the Long and Gray laboratories for helpful discussions. This work was supported by the NIH (DK124265 and DK130541 to J.Z.L. and DK137983 to MDB), the Phil & Penny Knight Initiative for Brain Resilience at the Wu Tsai Neurosciences Institute (research grant to J.Z.L.), the Ono Pharma Foundation (research grant to J.Z.L.). Graphical abstract was created with Biorender.com.

## AUTHOR CONTRIBUTIONS

Conceptualization, V.L.L., S.F., X.L., J.Z.L.; Investigation, V.L.L., S.F., L.W., X.L., W.W., X.S., S.D., J.L.B., U.A.T., C.A., A.C., B.H., L.M.R., D.F., T.Z., M.D.B., S.M.H.; Writing – Original Draft, V.L.L., S.F., L.W., T.Z., M.D.B., N.S.G., S.M.H., J.Z.L.; Writing – Review & Editing, V.L.L., S.F., L.W., T.Z., M.D.B., N.S.G., S.M.H., J.Z.L.; Resources, J.G.W., R.E.G., M.D.B., N.S.G., S.M.H., J.Z.L.; Supervision, J.G.W., R.E.G., T.Z., M.D.B., N.S.G., S.M.H., J.Z.L.

## STAR METHODS

### EXPERIMENTAL MODEL AND STUDY PARTICIPANT DETAILS

#### Animal information

All animal experiments were conducted in accordance with protocols approved by the Stanford University Administrative Panel on Laboratory Animal Care. Mice were maintained in a facility with 12-h light–dark cycles at 22 °C and approximately 50% relative humidity. The mice were fed a standard irradiated rodent chow diet, except where indicated, in which case a high-fat diet (D12492, Research Diets 60% kcal from fat) was used. Male C57BL/6J (stock number 000664) and male C57BL/6J DIO mice (stock number 380050) were purchased from the Jackson Laboratory. PTER-KO mice were obtained from heterozygous breeding pairs as described previously.^6^

## METHOD DETAILS

### Mouse PTER protein purification

The mouse PTER (mPTER) gene (Uniprot accession number Q60866) was codon-optimized for bacterial expression and synthesized as gBlocks by IDT. The gene fragment was cloned into the pET-20b vector with a C-terminal hexa-Histidine (His) tag and a N-terminal Strep tag. The pET-20b-mPTER plasmid was transformed into BL21 DE3 competent cells (ThermoScientific, EC0114) and grown in LB medium supplemented with ampicillin at 37 °C with shaking overnight. For protein expression, BL21 cells inoculated into autoinduction medium containing the following components: 10 g tryptone (FisherScientific, BP1421-500), 5 g yeast extract (FisherScientific, BP1422-500), 2 ml MgSO4 (1 M), 1 ml metal solution (0.05 M Feccir citrate, 0.02 M CaCl2, 0.02 M ZnSO4, 2 µM CoCl2, 2 µM CuSO4, 2 µM NiCl2, 2 µM Na2MoO4, 2 µM Boric acid), 20 ml salt solution (167.5g Na2HPO4, 85g KH2PO4, 53.4g NH4Cl and 17.8g Na2SO4 in 500 ml water in total) and 20 ml sugar solution (125 g glycerol, 12.5 g glucose and 50 g α-lactose in 500 ml water in total) in a total volume of 1 L. Cells were grown to optical density (OD600) of 0.5-0.7 at 37 °C, then incubated at15 °C overnight. Cells were harvested centrifugation at 8,000 rpm for 30 min at 4 °C and lysed in buffer (50 mM Tris-HCl, 135mM NaCl, pH7.4) by probe sonication on ice. Soluble fractions were obtained by centrifugation at 15,000 rpm for 30 min at 4 °C. The recombinant mPTER proteins were purfiied using Strep-Tactin resins (IBA, 2-1208-002), according to the manufacturer’s instructions. Bound mPTER proteins were eluted with buffer (2.5 mM D-Desthiobiotin, 25 mM Tris and 130 mM NaCl pH7.4). Purified proteins were aliquoted and stored at -80 °C for subsequent enzymatic assays.

### Expression screening of PTER orthologs in *E. coli*

Proteins were expressed in *E. coli* similar to what was previously described.^39^ Coding regions for human, sponge, fruitfly, rat, chicken, cat, zebrafish, mosquito, pig, frog, platypus, lamprey, hedgehog, armadillo, microbat, and Tasmanian devil were cloned by ligation independent cloning into expression vectors with the following elements: T7 polymerase promoter, N-terminal His6-SUMO tag, TEV protease cleavage site, and codon-optimized indicated species open reading frame. For protein expression in *E. coli* strain BL21 DE3, cells were grown to an optical density of ∼0.5, followed by induction of protein expression with 0.4 mM IPTG. Cultures were then incubated overnight at 18 °C before harvesting by centrifugation and freezing. For expression comparison, 75 ug of total bacterial lysate from each culture was mixed with SDS sample buffer, denatured by heating, separated on Novex™ Tris-Glycine Mini Protein Gels 4–20 % (Invitrogen), and transferred to nitrocellulose membranes. Relative expression levels of the recombinant proteins were assessed based on the intensity of anti-His bands.

### Sponge PTER protein expression and purification for crystallography

pET-T7-His6-SUMO-TEV-spongePTER was cloned as described above. Proteins were purified as previously described.^39^ For crystallography studies, sPTER was expressed in *E. coli* Rosetta 2(DE3)pLysS (Novagen). Cells were grown to an optical density of ∼0.5, followed by induction of protein expression with 0.4 mM IPTG. Cultures were then incubated overnight at 18 °C before harvesting by centrifugation and freezing.

For protein purification, cell pellets were thawed, resuspended in buffer D800 (20 mM HEPES, pH 7.5, 800 mM NaCl, 10 mM imidazole, 2 mM β-mercaptoethanol, 10% glycerol), and supplemented with protease inhibitors (aprotinin, leupeptin, and pepstatin). After sonication, soluble lysate was obtained by centrifugation for 30 min at 14,000 rpm (Thermo Scientific™ Sorvall X4R Pro-MD). Proteins were subsequently purified by Co^2+^ affinity chromatography and applied to an ion exchange column (5mL HiTrap Q HP). The flowthrough from ion exchange was concentrated and dialyzed overnight in dialysis buffer (25 mM Tris pH 7.5, NaCl 150 mM, 2 mM β-mercaptoethanol, 5 % glycerol). The dialyzed protein was subject to His6-SUMO tags cleavage by incubating with TEV protease for 2 h at room temperature. Protein was then adjusted to a final concentration of 50 mM imidazole to facilitate the remove of protease, cleaved tags, and uncleaved proteins on Ni^2+^ affinity chromatography (HisTrap HP, 5 ml); flowthrough containing tagless protein was collected and concentrated. Final purification was performed on a Superdex 200 column (10/300 GL, GE) equilibrated in gel filtration buffer (20 mM Tris-HCl, pH 8.5, 150 mM NaCl, 1 mM TCEP). Peak fractions were collected, concentrated by ultrafiltration, frozen in gel filtration buffer with 5% glycerol by volume, and stored at −80 °C until use. For crystallography experiments, no glycerol was added, and the protein was either used immediately or frozen at -80 °C in small, concentrated aliquots and used later.

### sPTER crystallization and structure determination

All crystals were obtained by the hanging-drop vapor-diffusion method at 16°C. Protein at 25mg/mL was mixed 1:1 (v/v) with well solution. For inhibitor-bound proteins, inhibitors at a concentration of 1.2 molar equivalents to the protein were added to the concentrated protein before crystallization. sPTER with inhibitors or products formed hexagonal crystals within 3 days. The crystallization buffer was 300mM MgCl_2_, 100mM Tris pH 8.5, and 20%PEG-8k. Apo sPTER crystals appeared within two days after microseeding with smaller crystals of the same morphology. The apo crystallization condition consisted of 200mM MgCl_2_, 100mM Tris pH 8.5, and 20%PEG-8k, and these crystals grew slowly relative to inhibitor/product-bound conditions and required microseeding to generate useable samples. Data were collected at the Stanford Synchrotron Radiation Lightsource (SSRL) beamline BL12-1.

Data reduction and integration were done using Xia-2 (Dials-Aimless pipeline).^40^ The best datasets displayed modest diffraction anisotropy, but the experimental electron density did not shown noticeable real-space distortions in any direction. Phase determination was achieved by using an AlphaFold2 Multimer model without metal ions^41^ as a search model and the Phaser MR module as implemented by CPP4.^42,43^ To account for modest anisotropy, conservative high resolution cutoffs were applied during structure refinement (Apo: 2.8Å, Products: 2.4Å, Mn(I/σ (I)) > 2.0). We specified one complex, a dimer, per asymmetric unit of the crystal. Models were improved by iterative rounds manual modification in Coot^44^ and refinement using REFMAC5.^45^ Waters and, for product-bound sPTER, acetate and taurine, were added at later stages of the refinement. Restraints for substrates were generated using Acedrg.^46^ Final models include residues 4-354 (product-bound) and 5-354 (apo). Crystallography statistics are shown in **Table S1**, and the coordinates have been deposited in the Protein Data Bank. Map and structural figures were generated with the PyMOL Molecular Graphics System, Version 2.5.8 Schrödinger, LLC.

### Molecular docking of PTERi and SAHA into sPTER

To determine the most likely poses of PTERi and SAHA, we used an unbiased diffusion-based docking routine, Diffdock.^23^ For this, we used our experimental apo-sPTER as a docking target and ran Diffdock using default parameters, requesting 10 models for each bound inhibitor. Docking results were consistent across runs and showed good concordance with poses determined using the experimental apo-PTER coordinates and AlphaFold2 coordinates as targets.^41^ We also examined docked PTERi and SAHA poses with respect to mouse PTER (AlphaFold2 model), and we observed similar results.

### hPTER and rPTER protein expression and purification

Human PTER (hPTER) was expressed utilizing the baculovirus expression vector system in Sf9 cells for hydrolysis assays due to low expression levels in *E. coli* BL21 strains. The hPTER was first cloned into a pFastBac backbone. The plasmid was then transformed into DH10 Bac *E. coli* cells containing the bacmid genome. Following transposition and antibiotic selection, bacmid DNA was extracted and used to infect Sf9 cells. After three rounds of baculovirus infection, the cells were harvested for subsequential protein purification. *E. coli* pellets expressing rPTER were obtained as described in the protein screening method section above. Frozen pellets were thawed in warm water and sonicated to obtain cell lysates. Lysates were incubated with cobalt resin on a rotator at 4°C for 1 hour. The cobalt resin was then pelleted, and the supernatant containing unbound proteins was discarded. Cobalt resin was resuspended and loaded onto a gravity column. After four washes to remove unbound proteins, proteins were eluted using 400mM imidazole. Subsequently, overnight dialysis was performed to remove the imidazole. Purified hPTER and rPTER were proceeded to N-acetyltaurine hydrolysis assays described below.

### N-acetyltaurine hydrolysis assays using recombinant PTER proteins

Enzymatic assays were performed using 200 ng of recombinant mPTER proteins incubated with 100 µM N-acetyltaurine (Cayman, 35169) and the indicated inhibitors in a 50 µl reaction buffer (50 mM Tris-HCl, 135 mM NaCl, pH 7.4) at 37 °C for 1 hour. The reactions were then quenched by the addition of 150 µl of a 2:1 (v/v) mixture of acetonitrile and methanol. The mixture was centrifuged at 15,000 rpm for 30 min at 4 °C to remove precipitated proteins and analyzed by LC-MS.

### HDAC activity assays using liver or recombinant proteins

Liver tissues were homogenized in HDAC reaction assay buffer (HEPES 50 mM, NaCl 137 mM, KCl 3 mM, MgCl2 1 mM, pH 8.0) at a ratio of 100 mg tissue per ml of buffer, and further sonicated. The lysates were centrifuged at 15,000 rpm for 30 min at 4 °C. The supernatant protein concentration was determined using a Nanodrop One. The HDAC assay was performed in 50 µl reaction volume containing the HDAC reaction assay buffer, 600 µg of the liver lysate, 100 µM HDAC substrate peptide Ac-Arg-Gly-Lys-Ac-Glu-AMC (custom synthesized by Elim Biopharm), and 10 µM of indicated inhibitors. The reaction mixture was incubated at 37°C for 1 hour, then quenched by the addition of 150 µl of a 2:1 (v/v) mixture of acetonitrile and methanol. The mixture was centrifuged at 15,000 rpm for 30 min at 4 °C to remove precipitated proteins and analyzed by LC-MS. Deacetylation activity assays using individual human HDAC and SIRT family members (HDAC 1-11 and SIRT 1, SIRT 2, SIRT 3, SIRT 4) were determined by Reaction Biology Corporation.

### Compound synthesis

Starting materials, reagents, and solvents were purchased from commercial suppliers and were used without further purification unless otherwise noted. All reactions were monitored using a Waters Acquity UPLC/MS system (Waters PDA eλ Detector, QDa Detector, Sample manager - FL, Binary Solvent Manager) using Acquity UPLC® BEH C18 column (2.1 x 50 mm, 1.7 μm particle size): solvent gradient = 85% A at 0 min, 1% A at 1.7 min; solvent A = 0.1% formic acid in Water; solvent B = 0.1% formic acid in Acetonitrile; flow rate: 0.6 mL/min. Reaction products were purified by flash column chromatography using CombiFlash®Rf with Teledyne Isco RediSep® normal-phase silica flash columns (4 g, 12 g, 24 g, 40 g or 80 g) and Waters HPLC system using SunFireTM Prep C18 column (19 x 100 mm, 5 μm particle size): solvent gradient = 80% A at 0 min, 10% A at 25 min; solvent A = 0.035% TFA in Water; solvent B = 0.035% TFA in MeOH; flow rate: 25 mL/min. ^1^H NMR spectra were recorded on 500 MHz Bruker Avance III spectrometers. Chemical shifts are reported in parts per million (ppm, δ) downfield from tetramethylsilane (TMS). Coupling constants (*J*) are reported in Hz. Spin multiplicities are described as *br* (broad), *s* (singlet), *d* (doublet), *t* (triplet), *q* (quartet), *m* (multiplet). Purities of assayed compounds were in all cases greater than 95%, as determined by reverse-phase LC-MS analysis. All synthetic methods and structural characterization for PTERi and PTERamide can be found in **Supplementary Information**.

### Drug treatment to mice

For intraperitoneal injections of mice with PTERi, the inhibitor was dissolved in a 3:1 mixture of Kolliphor:DMSO at a concentration of 100 mg/ml and injected into mice at 1 μl/g body weight for a final dose of 100 mg/kg body weight. Semaglutide (Cayman, catalogue number 29969) was dissolved in saline at a concentration of 60 nM or 400 nM and injected into mice at 5 μl/g body weight for a final dose of 0.3 or 2 nmol/kg body weight, respectively.

### Conditioned taste aversion tests

Mice were single housed and acclimated (up to 3 days) to drinking from two water bottles to confirm lack of side preference prior to habituation. Mice were then habituated to overnight water restriction (days 1-3) followed by 1 h water bottle presentation (two bottles) and saline injection (IP). On day 4 to begin conditioning, mice were instead given a novel 0.15% saccharin solution in both bottles instead of water for 30 minutes, followed by a injection of either saline, PTERi (100 mg/kg, IP), or the positive control LiCl (225 mM, 0.1mL per 10g of body, IP). Access to saccharin water was allowed for an additional 30 min and was then changed back to water until the next restriction. Day 5 was used as a washout period using the days 1-3 water bottle protocol. A second conditioning period was performed on day 6 followed by a washout period on day 7. On day 8, a standard two bottle preference test (saccharin versus water) was used to assess CTA development to the saccharin solution (1 h presentation after overnight water restriction). Fluid intake was measured for both saccharin and water.

### Effect of anti-GFRAL neutralizing antibody on PTERi-induced feeding suppression

DIO mice were injected with GFRAL neutralizing antibody or control IgG antibody (subcutaneous, 10 mg/kg in PBS). The next day, mice were separated into individual cages and fasted for 6 hours. At 7 PM, mice were injected with either vehicle or PTERi (IP, 100 mg/kg) and provided with high fat diet. Food intake 3 hours post-injection was measured.

### Preparation of mouse blood for LC-MS analysis

Mouse blood was collected in heparin tubes and centrifuged at 5000 rpm for 5 min at 4°C to obtain plasma. A 50 µl aliquot of plasma was mixed with 150 µl of a 2:1 (v/v) mixture of acetonitrile and methanol and vortexed for 30 s to precipitate proteins. The mixture was then centrifuged at 15,000 rpm for 30 min at 4 °C and the supernatant was transferred to a LC–MS vial for subsequent analysis. For time course experiments, approximately 100 µl of blood was collected at each time point. A 20 µl aliquot of plasma was used for metabolite isolation, and the volume of the acetonitrile:methanol mixture was proportionally reduced to maintain the 1:3 (plasma:solvent) ratio.

### Western blotting

For mouse tissue samples, tissues were dissected, weighed, immediately frozen on dry ice and stored at −80 °C. Tissues were homogenized in 0.5 ml of cold RIPA buffer (containing Deacetylase Inhibitor Cocktail, MedChemExpress, HY-K0030) using a Benchmark BeadBlaster Homogenizer at 4 °C. The homogenate was further sonicated and quantified using a tabletop Nanodrop One and analyzed by western blot. Proteins were separated on Novex™ Tris-Glycine Mini Protein Gels 4–20 % (Invitrogen) and transferred to nitrocellulose membranes. Equal loading was confirmed by Ponceau S staining. Membranes were blocked with Odyssey blocking buffer for 30 min at room temperature and incubated overnight at 4 °C with primary antibodies: 1:5000 dilution rabbit anti-β-actin antibody (Abcam, ab8227), 1:1000 dilution rabbit anti-6xHis antibody (Abcam, ab9108), 1:1000 anti-Histone H4K16Ac antibody (Abcam, ab109463), 1:1000 anti-Histone H3K27ac antibody (Active Motif, 39134), 1:1000 anti-oxphos (Thermo 45-8099). Membranes were washed three times with PBST (0.05% Tween-20 in PBS) and stained with species-matched secondary antibodies (1:10000 dilution goat anti-rabbit IRDye 800RD (LI-COR, 925-68070) and 1:10000 dilution goat anti-mouse IRDye 680RD (LI-COR, 925-68070)) at room temperature for 1 h. Blots were further washed three times with PBST and imaged with the Odyssey CLx Imaging System.

### Measurement of mRNA levels by qPCR

Total RNA was isolated using RNAeasy mini kit (Qiagen) according to the manufacturer instructions. cDNA was reverse transcribed using the High-capacity cDNA reverse transcription kit (Thermo). The primers used for qPCR are shown in the **Key Resources Table**.

### Measurements of metabolites by LC-MS

Metabolite measurements were performed using an Agilent 6545 Quadrupole time-of-flight LC–MS instrument as previously described.^6^ MS analysis was conducted using electrospray ionization (ESI) in negative mode. The dual ESI source parameters were configured as follows: the gas temperature, 250 °C; drying gas flow,12 l/min; nebulizer pressure, 20 psi; capillary voltage, 3,500 V; and the fragmentor voltage, 100 V. The separation of polar metabolites was achieved using a Luna 5 μm NH2 100 Å LC column (Phenomenex 00B-4378-E0) with normal phase chromatography. Mobile phases were as follows: buffer A, 95:5 water:acetonitrile with 0.2% ammonium hydroxide and 10 mM ammonium acetate; buffer B, acetonitrile. For the in vitro assay, the LC gradient initiated at 100% B with a flow rate of 0.7 ml/min from 0 to 0.2 min. The gradient was then linearly increased to 20% A/80% B at a flow rate of 0.7 ml/min from 0.4 to 1.1 min. From 1.1 to 4 min, the gradient was linearly increased to 50% A/50% B at the same flow rate. The gradient was maintained at 50% A/50% B at a flow rate of 0.7 ml/min from 4min to 4.5min. The gradient was decreased to 0%A/100%B from 4.5min to 5min and maintained at 0%A/100%B from 5min to 6min. N-acetyltaurine (Cayman, 35169) eluted around 4 min and taurine (Sigma, T0625-500G) eluted around 4.1 min under the above conditions. For in vivo assay samples, the LC gradient initiated at 100% B with a flow rate of 0.7 ml/min from 0 to 2 min. The gradient was then linearly increased to 50% A/50% B at a flow rate of 0.7 ml/min from 2 to 20 min. From 20 to 25 min, the gradient was maintained at 50% A/50% B at a flow rate of 0.7 ml/min. The gradient was maintained at 0%A/100%B from 25 min to 30 min. N-acetyltaurine (Cayman, 35169) eluted around 11.5 min and taurine (Sigma, T0625-500G) eluted around 12.8 min under the above conditions.

## QUANTIFICATION AND STATISTICAL ANALYSIS

All data are expressed as mean ± SEM as indicated in the legends. A student’s two-tailed unpaired t-test was used for pair-wise comparisons. For comparisons of multiple groups, P-values were calculated by two-way ANOVA. Unless otherwise specified, statistical significance was set as P < 0.05. P < 0.05 was used to determine significance.

