## Supplemental Figures and Table S1 for "A small molecule PTER-selective inhibitor reduces food intake and body weight"

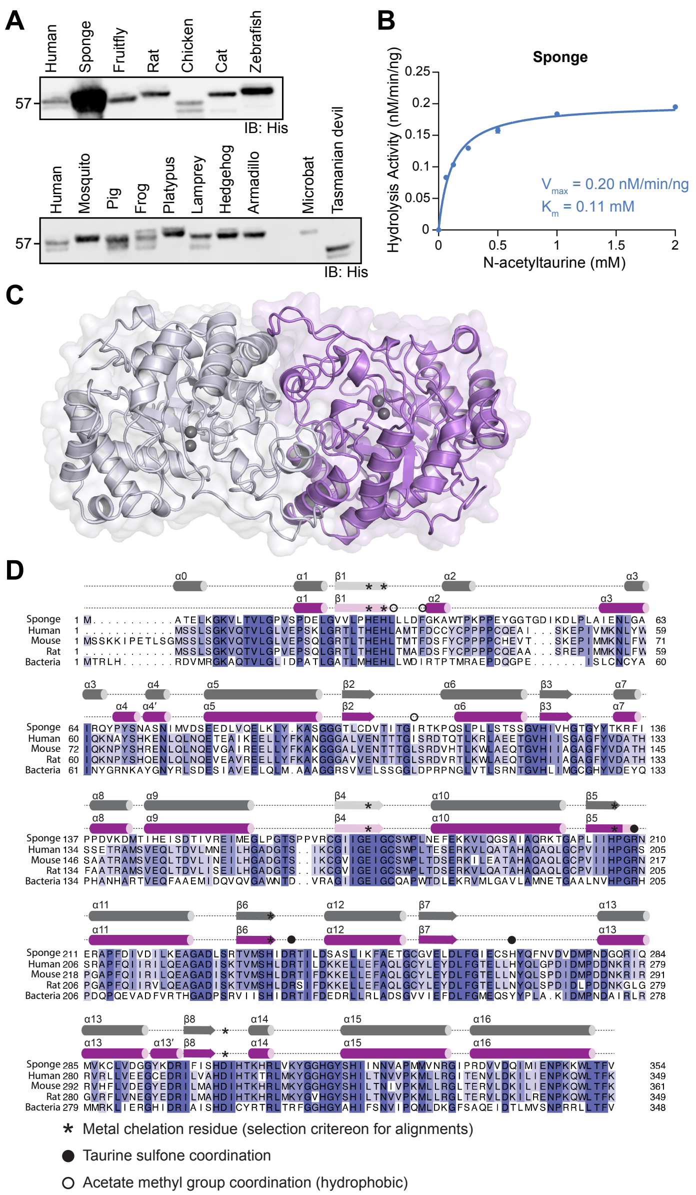


**Fig. S1. Biochemical characterization of sponge PTER and crystal structure and sequence alignment of apo sponge PTER.**

**(A)** Anti-His Western blot showing relative expression levels of recombinant His6-Sumo-PTER homologs in *E. coli*. 75 μg total bacterial lysate induced to express the indicated PTER homolog was loaded per lane.

**(B)** N-acetyltaurine hydrolysis activity following incubation of 200 ng purified recombinant sponge PTER with the indicated concentrations of N-acetyltaurine. Data are presented as the mean ± SEM and were fitted to Michaelis–Menten kinetics (solid line) using GraphPad Prism.

**(C)** Crystal structure of apo-sPTER homodimer. The structure is shown as a cartoon with a semi-transparent surface (grey, chain A; purple, chain B) with zinc (Zn) ions shown as grey spheres.

**(D)** Sequence alignment of sponge, human, mouse, rat PTERs and bacterial PTE, colored by amino acid conservation. Predicted secondary structures are displayed above the alignment (grey, PTE; purple, PTER), with major interface residues annotated as the key below.

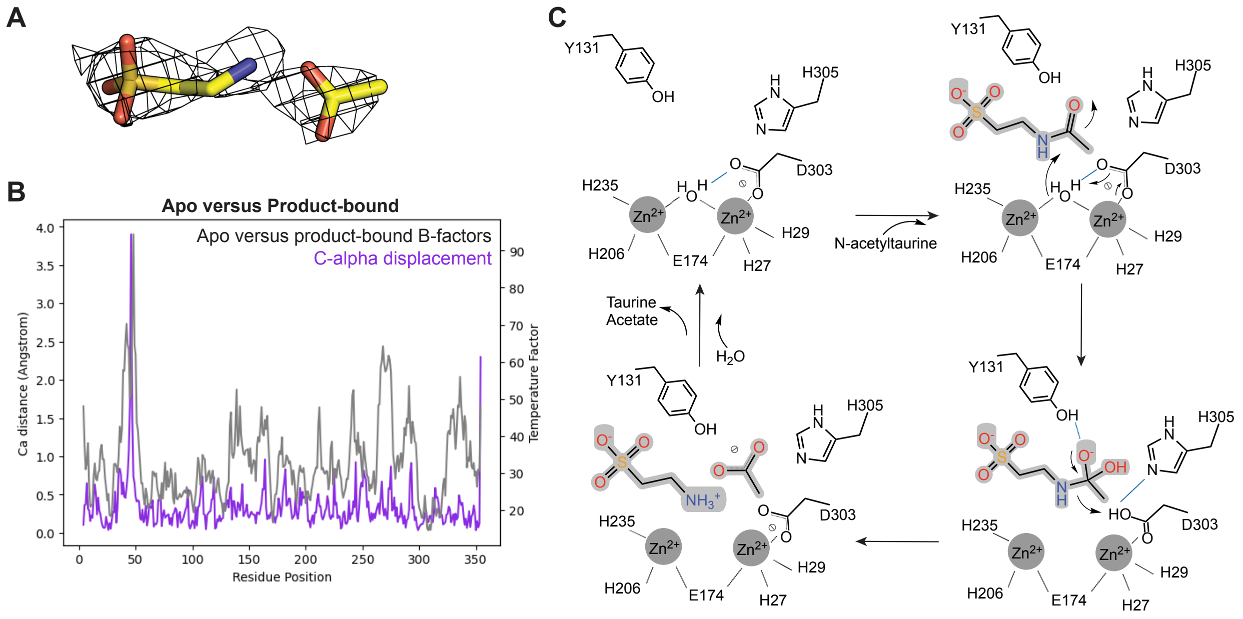


**Fig. S2. Additional analysis of sPTER in complex with natural acetate and taurine ligands.**
**(A)** 2Fo–Fc electron density map at 1.0 σ for taurine and acetate in the product-bound PTER co-crystal structure.
**(B)** Plot of carbon-alpha (Cα) displacement and temperature factors for sPTER upon binding products acetate and taurine. Large Cα displacements correlate to high temperature factors.
**(C)** Proposed catalytic mechanism of N-acetyltaurine deacetylation by PTER.

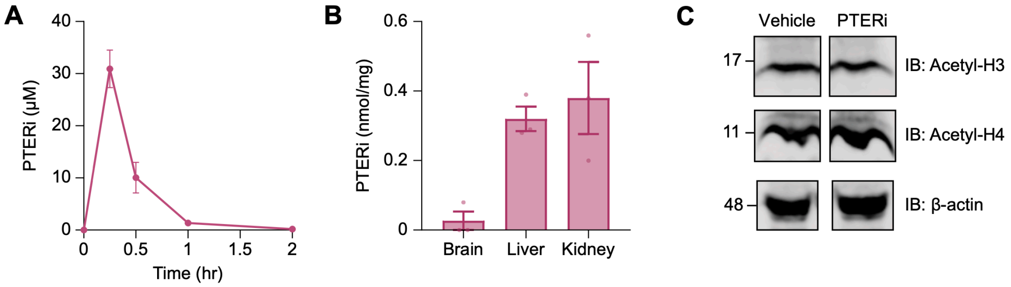


**Fig. S3. Additional characterization of acute PTERi effects in mice.**
**(A-C)** Plasma drug levels **(A)**, tissue drug levels **(B)**, and liver histone acetylation levels **(C)** after a single administration of vehicle or PTERi (100 mg/kg, IP) to 8-week old **(A,B)** or 12-week old **(C)** male lean mice. For **(A,B)** N=3 mice/group. For **(C)**, tissues were harvested at 1 h after injection.
All data are shown as mean ± SEM.

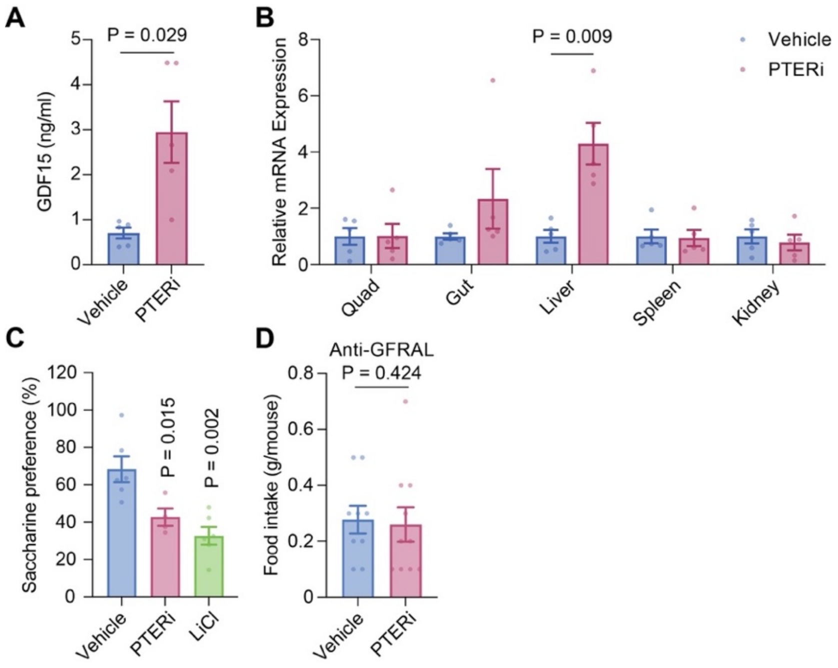


**Fig. S4. Role of GDF15/GFRAL in the anorexigenic effects of PTERi in mice.**
**(A,B)** Plasma GDF15 (**A**) and tissue *Gdf15* mRNA (**B**) levels 1 hr after a single administration of vehicle or PTERi (100 mg/kg, IP) to 18-week-old male DIO mice. N=5 mice/group.
**(C)** Saccharin preference (as percentage of total fluid consumption) during a conditioned taste aversion test following vehicle, PTERi (100 mg/kg, IP) or LiCl treatment (100 mg/kg, IP) to 7-8 week-old male lean mice. N=6 mice for vehicle and LiCl, N=4 mice for PTERi.
**(D)** Food intake over 3 h after a single administration of vehicle or PTERi (100 mg/kg, IP) to 14-week-old male DIO mice that had been treated with a neutralizing anti-GFRAL antibody (10 mg/kg, subcutaneous) one day prior. Initial body weights, mean ± SEM: vehicle, 38.4 ± 1.3 g; PTERi, 38.2 ± 1.1 g.

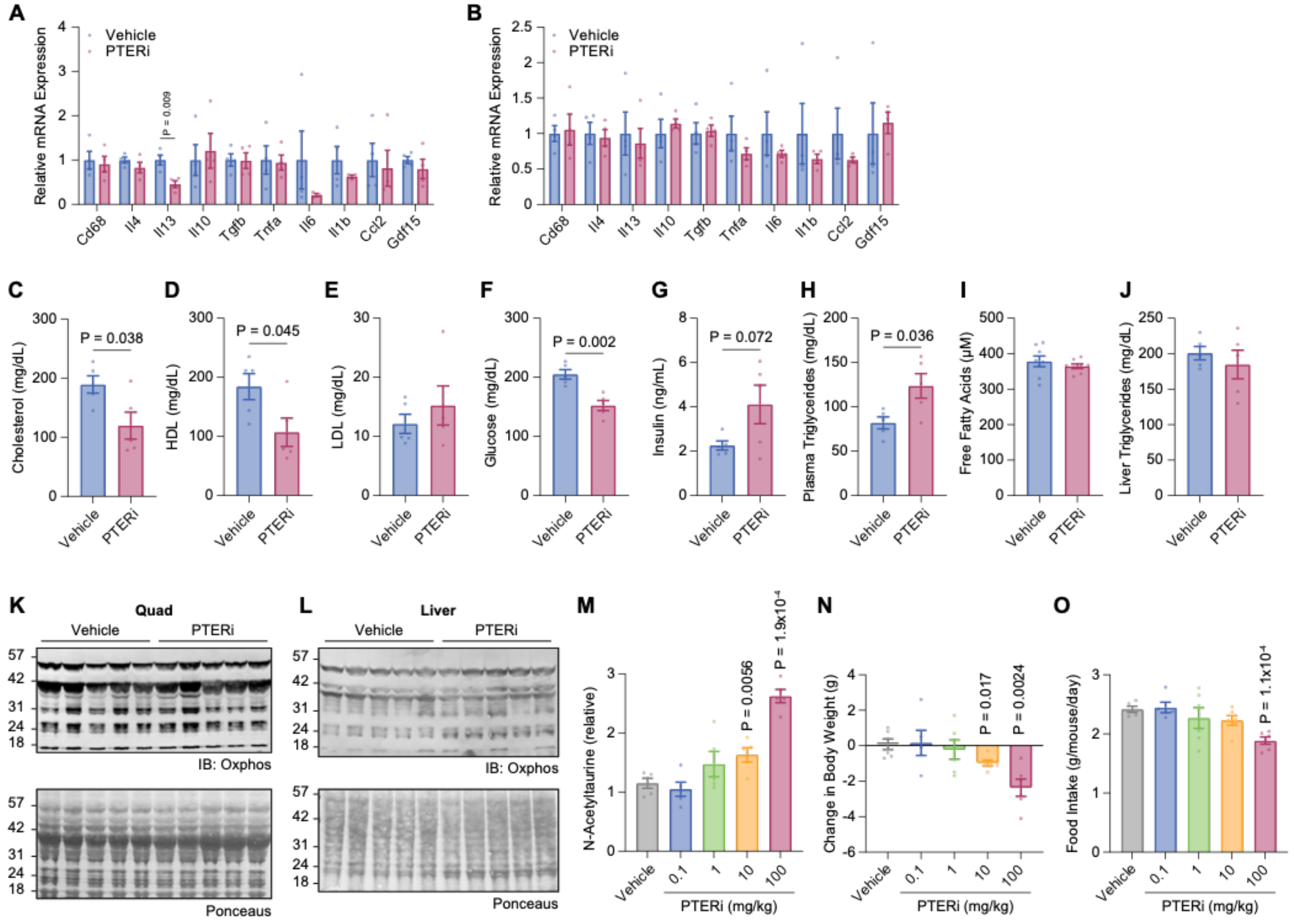


**Fig. S5. Additional characterization of chronic PTERi administration in DIO mice.**

**(A-L)** Epididymal white adipose tissue (eWAT) gene expression **(A)**, liver gene expression **(B)**, plasma cholesterol **(C)**, HDL **(D)**, LDL **(E)**, glucose **(F)**, insulin **(G)**, triglycerides **(H)**, free fatty acids **(I)**, liver triglycerides **(J)**, and anti-oxphos Western blotting of muscle tissues **(K)** or anti-oxphos Western blotting of liver tissues **(L)** from 18 week-old male DIO mice following chronic administration with vehicle or PTERi (100 mg/kg/day, IP) for 27 days. Tissues were harvested 24 h after the last dose of PTERi. Initial body weights, mean ± SEM: vehicle, 39.3 ± 1.0 g; PTERi, 38.1 ± 0.9 g, P > 0.05.

**(M)** Relative plasma N-acetyltaurine levels after a single administration of vehicle or PTERi at the indicated doses (0.1-100 mg/kg, IP) to 18-week-old male DIO mice. N=5 mice/group.

**(N,O)** Change in body weight **(N)** and cumulative food intake **(O)** of 18-week-old male DIO mice following chronic administration with either vehicle, or PTERi (0.1-100 mg/kg, IP). For vehicle, 1 mg/kg, 10 mg/kg, 100 mg/kg PTERi, N=6 mice/group. For 0.1 mg/kg PTERi, N=5 mice. Initial body weights, mean ± SEM: vehicle, 41.3 ± 2.2 g; 0.1 mg/kg PTERi, 41.5 ± 1.5 g; 1 mg/kg PTERi, 41.4 ± 2.0 g; 10 mg/kg PTERi, 42.0 ± 0.9 g; 100 mg/kg PTERi, 40.5 ± 1.9 g.

All data are shown as mean ± SEM. All P-values were calculated by Student’s *t*-test.

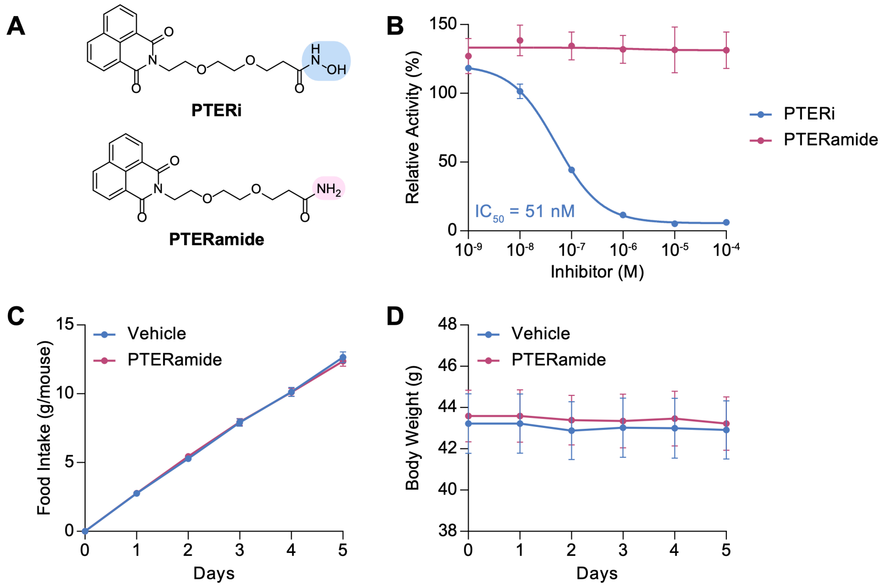


**Fig. S6. In vitro and in vivo characterization of the inactive analog PTERamide.**
**(A)** Chemical structures of PTERi (top) and PTERamide (bottom).
**(B)** Dose-response curves of PTERi (blue) and PTERamide (red) in N-acetyltaurine hydrolysis activity assay. 200 ng of purified recombinant mPTER was incubated with 100 μM N-acetyltaurine and the indicated concentration of inhibitor for 1 h at 37ºC.
**(C,D)** Cumulative food intake (**C**) and change in body weight (**D**) of 18 week-old male DIO mice following chronic administration of either vehicle or PTERamide (100 mg/kg/day, IP).
All data are shown as mean ± SEM. $IC_{50}$ values for **(B)** was determined from the dose-response curves via nonlinear regression analysis using GraphPad Prism. For **(C,D)**, N=12 mice per group.

| | Apo sPTER | Tau Ace sPTER |
|---|---|---|
| **Data collection** | | |
| Resolution (Å) | 31.33–2.75 | 42.33–2.4 |
| Wavelength (Å) | 0.97946 | 0.97946 |
| Space group | $P2_1$ | $P2_1$ |
| Unit cell dimensions (a, b, c) (Å) | 82.85, 48.37, 83.23 | 83.99, 48.95, 85.27 |
| Unit cell angles (α, β, γ) (°) | 90.00, 96.98, 90.00 | 90.00, 98.56, 90.00 |
| Molecules per ASU | 2 | 2 |
| Total reflections | 75876 | 143230 |
| Unique reflections | 16884 | 25850 |
| Completeness (%)[a] | 97.4 (95.9) | 95.4 (97.1) |
| Multipilicity[a] | 4.5 (4.4) | 5.5 (5.6) |
| I/σ[a] | 6.0 (2.7) | 6.4 (1.4) |
| Rmerge(I)[a] | 0.202 (0.692) | 0.203 (1.579) |
| CC1/2(%)[a] | 88.1 (58.1) | 95.4 (33.7) |
| **Refinement** | | |
| Resolution (Å) | 31.12–2.8 | 42.33-2.4 |
| All reflections | 16498 | 27248 |
| free | 798 | 1436 |
| Rwork (%) | 20.73 | 18.22 |
| Rfree (%) | 28.12 | 27.59 |
| Wilson B-factor ($Å^2$) | 24.9 | 31.4 |
| protein | 29.34 | 38.92 |
| zinc | 20.47 | 36.19 |
| ligand | - | 91.51 |
| water | 19.12 | 37.59 |
| **Structure Statistics** | | |
| Number of atoms (protein dimer) | | |
| protein | 5434 | 5457 |
| zinc | 4 | 4 |
| ligand | - | 22 |
| water | 2 | 281 |
| r.m.s.d. bond distances | 0.0061 | 0.0059 |
| r.m.s.d. bond angles | 1.5969 | 1.5426 |
| Rotamer outliners (%) | 7 | 4 |
| Ramachandran plot (%) | | |
| favored | 91 | 95 |
| allowed | 9 | 5 |
| outliers | 2 | 1 |

**Table S1. Crystallography statistics of apo- and product-bound sPTER structures.** Superscript a, highest resolution shell values in parenthesis.
